# Competitive Microtubule Binding of PEX14 Coordinates Peroxisomal Protein Import and Motility

**DOI:** 10.1101/2020.07.23.218024

**Authors:** Maren Reuter, Hamed Kooshapur, Jeff-Gordian Suda, Alexander Neuhaus, Lena Brühl, Pratima Bharti, Martin Jung, Wolfgang Schliebs, Michael Sattler, Ralf Erdmann

## Abstract

PEX14 functions as peroxisomal docking protein for the import receptor PEX5. For docking, the conserved N-terminal domain of PEX14 (PEX14-NTD) binds amphipathic alpha-helical ligands, typically comprising one or two aromatic residues, of which human PEX5 possesses eight. Here, we show that the PEX14-NTD also binds to microtubular filaments *in vitro* with a dissociation constant in nanomolar range. PEX14 interacts with two motifs in the C-terminal region of human ß-tubulin. At least one of the binding motifs is in spatial proximity to the binding site of microtubules (MT) for kinesin. Both PEX14 and kinesin can bind to MT simultaneously. Notably, binding of PEX14 to tubulin can be prevented by its association with PEX5. The data suggest that PEX5 competes peroxisome anchoring to MT by occupying the ß-tubulin-binding site of PEX14. The competitive correlation of matrix protein import and motility may facilitate the homogeneous dispersion of peroxisomes in mammalian cells.

## Introduction

Peroxisomes are ubiquitous single membrane bound cell organelles with a remarkable functional and morphological plasticity, depending on the cell or tissue type, organism and prevailing environmental conditions [1]. Peroxisomes can multiply by *de novo* formation from the ER and mitochondria and by growth and division of pre-existing ones [2, 3]. The biogenesis and dispersal of newly formed organelles, inheritance and autophagic degradation involves association with the cytoskeleton, either microtubules (MT) or actin filaments [4]. While association of peroxisomes with actin bundles is accompanied by oscillatory (vibrational) motions, the bi-directional MT-dependent movements are characterized by longer distances (>2 μm), higher speed (>0.2 μm/s) and lower frequency (<10%) [5, 6]. As shown for transport of other cellular organelles in mammalian cells (mitochondria and lysosomes), motor proteins of the dynactin-dynein or kinesin family pull peroxisomes along microtubule tracks [4, 7]. Whereas several mechanistic studies on motor-driven peroxisome transport have been published [7, 8], not much is known about the initiation and coordination of the transport processes. Within the last years, several peroxisome-associated proteins have been identified that mediate interaction with motor proteins, adaptor proteins or other constituents of the cytoskeleton. Very recently, the well-studied motor adaptor mitochondrial Rho GTPase 1 (MIRO1) has been shown to associate with peroxisomal membranes [9, 10]. As demonstrated for mitochondria previously, peroxisome-targeted MIRO1 can recruit microtubule-associated molecular motors by engaging the kinesin adaptors TRAK1/2 and thus, initiate fast, long-bidirectional transport of peroxisomes along MT [9, 10]. Based on a two-hybrid screen, PEX1 was identified as a potential peroxisomal membrane anchor for the motor protein KifC3, a member of the kinesin-14 family [11]. PEX1 is a peroxisomal biogenesis factor required for peroxisomal import machinery for matrix proteins.

Interestingly, the peroxisomal biogenesis factor PEX14 is involved in peroxisome positioning via MT [12]. This peroxin has been originally identified in *Saccharomyces cerevisiae* as a peroxisomal docking protein for cargo-loaded import receptors, PEX5 and PEX7, and was therefore assigned as the “point-of-convergence” of both import pathways [13, 14]. Furthermore, yeast PEX14 together with the peroxisomal matrix protein receptor PEX5 forms a transient ion conducting channel that can expand in diameter and is supposed to translocate matrix proteins across the peroxisomal membrane [15]. In addition, PEX14 is required for pexophagy, the selective degradation of peroxisomes [16–18]. In mammalian cells, the autophagic removal of superfluous peroxisomes under starvation is initiated by PEX14-mediated recruitment of MT-associated protein 1A/1B light chain 3 (LC3) [19, 20].

In addition to the functions listed above, human PEX14 is also required for MT-based peroxisome morphogenesis and movement. This function was first attributed to PEX14 by the observation that the typical MT-dependent movements of “large” peroxisomes is affected in PEX14-deficient fibroblasts from Zellweger patients [12]. Furthermore, the accumulation of peroxisomes at the cell periphery due to overexpression of certain MIRO1-constructs is not seen in PEX14-deficient cells, suggesting that PEX14 is indispensable for MIRO1-mediated peroxisomal movement [10]. Equally, the typical MIRO1-induced elongation of dividing peroxisomes and alignment of peroxisomes along the mitotic spindle is affected in the absence of PEX14 [9, 21]. In addition to the physiological evidence for a role of PEX14 in peroxisome motility and peroxisome movement along microtubules, physical association of purified PEX14 fragments with MT was demonstrated by pull-down assays and binding of purified PEX14 to the cytoskeleton of fixed cells [12, 22]. Notably, the MT-binding site of PEX14 was mapped to the well-conserved N-terminal domain of PEX14 (PEX14-NTD), which also interacts with multiple sites on PEX5 and thus contributes to receptor docking and pore formation. The eight high-affinity PEX14-binding sites of human PEX5 are amphipathic alpha-helical pentapeptides that are characterized by large hydrophobic residues at both ends, either tryptophan or leucine/valine at position 1 and the aromatic residues phenylalanine or tyrosine at position 5 [23, 24]. Moreover, the PEX14-NTD interacts with PEX19, the import receptor for peroxisomal membrane proteins [23]. Based on the diversity of binding partners of PEX14-NTD, a correlation between protein import into peroxisomes and microtubular anchoring of peroxisomes has been suggested [4].

Here, we investigated this correlation and analyzed the *in vivo* and *in vitro* interaction of PEX14-NTD with its binding partners. *In vitro* binding assays and NMR studies revealed that both PEX5 and ß-tubulin bind to PEX14-NTD. We show that the PEX14-NTD interacts with the C-terminal region of ß-tubulin that is also essential for binding of motor proteins. Interestingly, the PEX14-NTD binding sites for PEX5 and ß-tubulin overlap and both proteins compete for PEX14-NTD binding. Accordingly, overproduction of PEX5 in living cells results in shorter moving distance and lower speed of motile MT-associated peroxisomes. The structural and functional features suggest that peroxisomal PEX14 that is not occupied by PEX5 functions as a tethering factor for microtubules, which enables the binding of motor proteins and movement of peroxisomes for an efficient dispersion of peroxisomes throughout the cell.

## Results

### Competitive binding of PEX5 and MT to PEX14

The *in vitro* interaction of PEX5, PEX14 and MT was analyzed in a newly established ELISA-based assay. PEX5 was coated onto microtiter plates and bound monomeric His-PEX14-NTD or dimeric GST-PEX14-NTD was quantified (**Fig. 1**). In agreement with previous studies [23–25], PEX5 binds to GST- and His-tagged versions of PEX14-NTD with dissociation constant (*K_D_*) values of about 0.5 nM and 1.3 nM, respectively (**Fig. 1A**). To analyze MT association of PEX5 and PEX14, the plates were coated with purified fractions of porcine tubulin (**Supplementary Figure 1**). While His-PEX5 showed no association with MT (**Supplementary Figure 2A**), both PEX14-NTD constructs interacted with tubulin (**Fig. 1B**). Consistent with previous observations [22], the GST-tagged dimeric PEX14-NTD binds with higher affinity to MT than the monomeric protein (**Fig. 1B**). The binding affinity for the formation of GST-PEX14-NTD/MT complex corresponds to a *K_D_* in the low nanomolar range (0.7–2.3 nM) in five independent experiments (**Fig. 1B, Fig. 3B and Fig. 7C, D**), while the binding affinity for the monomeric His-PEX14-NTD/MT interaction is at least two orders of magnitude weaker. It should be noted that saturation of His-PEX14 binding could not be achieved under these experimental conditions. Since GST alone shows non-specific binding to MT at concentrations higher than 1 μM (**Supplementary Figure 2B**), it is not clear whether the high MT affinity of GST-PEX14-NTD is caused by the dimeric appearance of PEX14, as suggested earlier [22], or by additional binding of the GST fusion tag.

**Figure 1.**
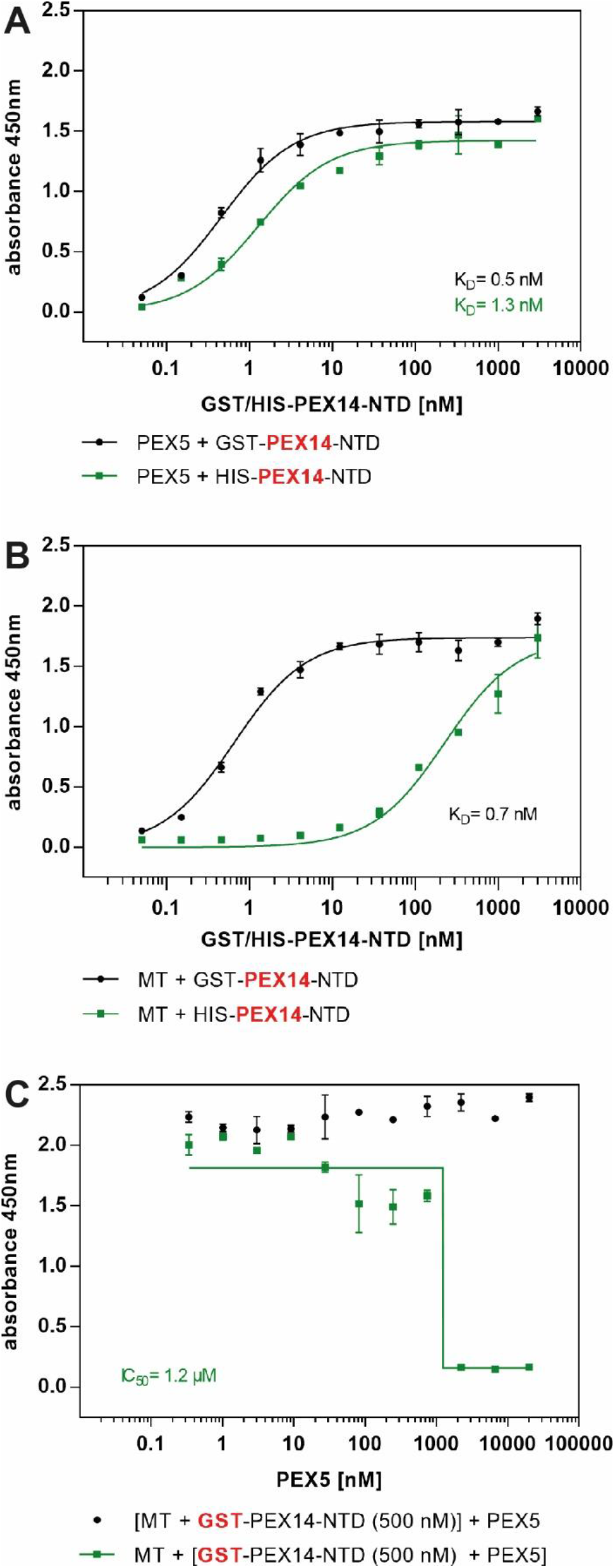
High-affinity binding of PEX14-NTD to MT can be blocked by PEX5. Microtiter plates were coated either with 0.01 mg/ml of purified PEX5 (A) or MT (B) and incubated with various concentrations (0.05 nM to 3 μM) of GST (black)- or His (green)-tagged PEX14-NTD. For the analysis of competitive effects of PEX5 on the MT/PEX14-NTD complex formation, MT-coated plates were first loaded with 500 nM GST-PEX14-NTD and then with varying concentrations of PEX5 (C, black), or the coated plates were incubated with a pre-incubated mixture of 500 nM GST-PEX14-NTD and varying concentrations (0.3 nM up to 20 μM) of PEX5 (C, green). Bound proteins or fragments which were immunodetected by primary specific antibodies (anti-PEX14 (A, B) and anti-GST (C)) are indicated in figure legends in red. The experimental data were fitted to a standard one-site model using non-linear regression and shown with standard errors (n=2).

To analyze whether full-length PEX5 has an effect on the PEX14/MT complex formation as previously proposed [12], MT-coated plates were first decorated with 500 nM GST-PEX14-NTD and then incubated with increasing concentrations of PEX5 (**Fig. 1C**, black). Under these conditions, no PEX14 was released, indicating a very low dissociation rate of the PEX14/MT complex. However, preincubation of GST-PEX14-NTD (500 nM) with increasing concentrations of PEX5 prior to incubation with MT-coated plates resulted in a significant block of the binding (**Fig. 1C**, green). The IC50 values of around 1.2 μM suggest that binding of PEX14 blocks its binding to MT. These data suggest that MT and PEX5 compete for the same binding site on PEX14.

To investigate the physiological relevance for this competition, peroxisome motility upon overexpression of PEX5 was monitored in living cells. To this end, human fibroblasts were transfected with a bicistronic vector expressing PEX5 as well as the fusion protein EGFP-PEX26 as a peroxisomal marker. This resulted in a significant overexpression of PEX5 as indicated by the corresponding steady state level (**Fig. 2A**). Motility of peroxisomes was studied using time-lapse imaging in the PEX5 overproducing and non-transfected human cells as control. Projections of fast directional and saltatory movement of labelled peroxisomes were recorded every six seconds and analyzed for run length and velocities. As shown in **Fig. 2B**, overexpression of PEX5 resulted in travelling distances of 2.15 μm on average before stalling of the organelles. In comparison, peroxisomes in normal environment travel around two-fold longer distances without interruption. In addition, overexpression of PEX5 reduces the average velocity of peroxisomes from 0.37 to 0.24 μm/s (**Fig. 2B**). It is noteworthy that the lower speed value lies within a range where fast actin- and slow MT-dependent moves are difficult to distinguish [5, 6]. In line with *in vitro* binding results (**Fig. 1**), our data suggest that PEX5 overexpression in living cells affects microtubule-dependent motility of peroxisomes by physically blocking the MT-binding site of PEX14.

**Figure 2:**
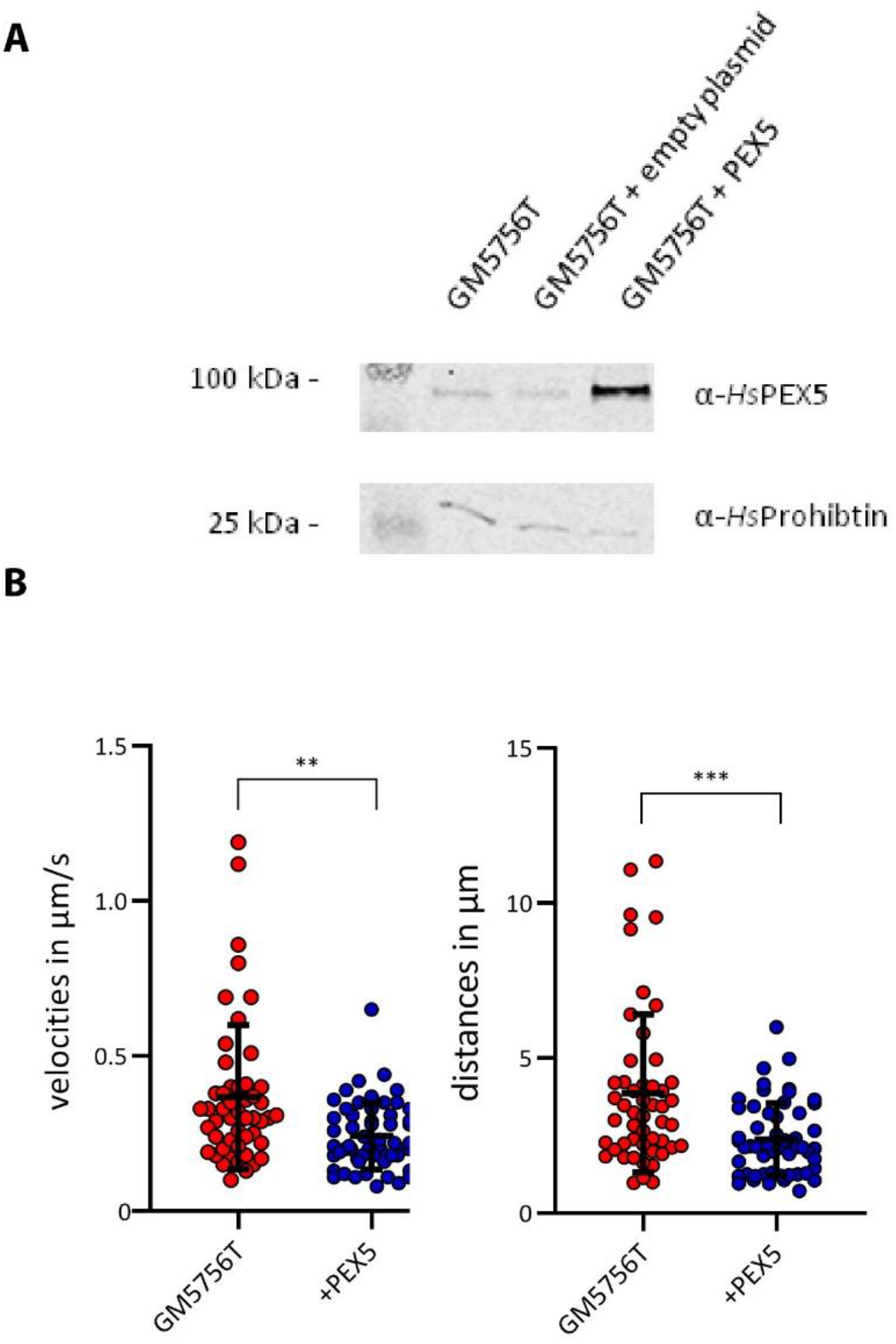
Overexpression of PEX5 affects peroxisome motility in human fibroblasts. GM5756T human fibroblasts were transfected with a bicistronic vector encoding EGFP-PEX26 together with PEX5 or without PEX5 (empty plasmid). Lysates were separated by SDS-PAGE and analyzed by immunoblotting using anti *Hs*PEX5 or anti *Hs*Prohibitin antibodies to demonstrate equal sample loading (A). The transfected fibroblasts expressing PEX5 under control of a CMV promoter displayed at least 10-fold overproduction of the PTS1 receptor. (B) MT-dependent motility of peroxisomes was studied in PEX5-overexpressing fibroblasts by life-cell imaging in at least 100 cells. The statistical analysis of all analyzed bidirectional saltations in GM5756T cells expressing PEX5 at endogenous (red dots) or enhanced (blue dots) differ in in terms of run length (distances) and velocity. (C) Overexpression of PEX5 results in significant reduction of velocity (p>0.01, **) and reveal shorter run distances (p>0.001, ***). Error bars represent SEM. Statistics were analyzed using non-parametric Mann-Whitney-U test.

### PEX5-interacting surface residues of PEX14-NTD are critical for MT-binding

To investigate if the binding sites of PEX14 for PEX5 and MT overlap, we applied structure-based mutagenesis of PEX14 residues that have previously been identified as critical for receptor binding [23, 24, 26]. K56 forms a salt bridge with negatively charged surface residues of PEX5 while F52 forms part of the hydrophobic PEX5-binding groove of PEX14 (**Fig. 3A**). These residues, as well as K54 as negative control, were replaced by alanine and the MT-binding capacity of the resulting PEX14 variants was determined using ELISA binding assays. For wild-type PEX14, the apparent *K_D_* of binding to MT is 2.1 +/- 0.2 nM and the F52A and K56A substitutions result in a significant increase of the *K_D_* to 39 +/- 7.4 nM and 12.5 +/- 1.3 nM, respectively (**Fig. 3B**). Combination of the F52A mutation with K56A resulted in an even more drastic effect; the double mutant PEX14 binds to MT with a *K_D_* of 89 +/- 17 nM (**Fig. 3C**), thus about 30-fold higher than the interaction with the wild-type protein. By contrast, the PEX14 K54A variant binds to MT with a *K_D_* of 3.9 +/- 0.2 nM similar to the wild-type protein. These data indicate that the PEX14 residues F52 and K56 that are critical for PEX14 binding of PEX5 are critical for PEX14 binding to MT, suggesting that the binding mode of PEX14 to both PEX5 and MT might be similar.

**Figure 3.**
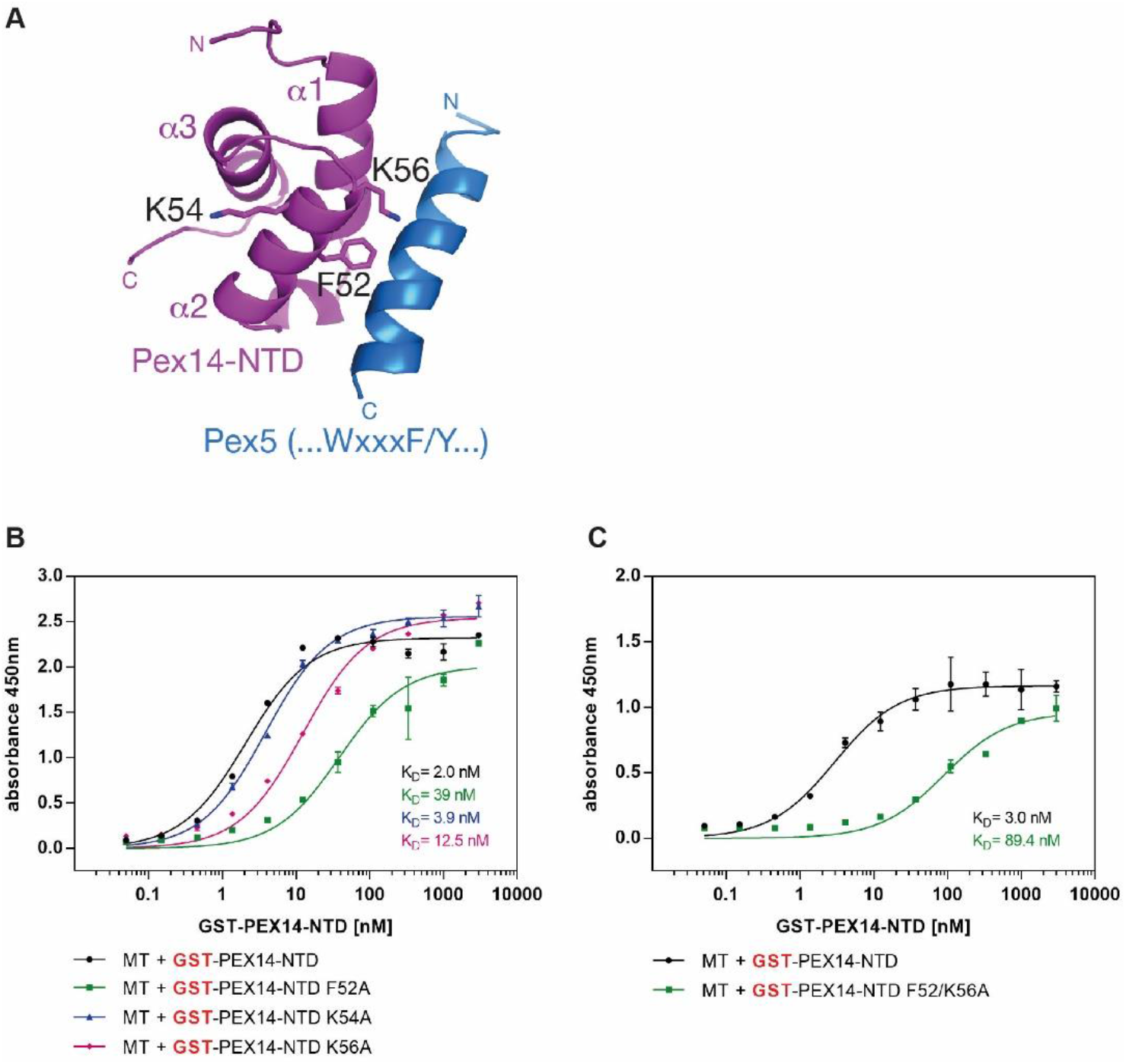
Identification of critical residues of the PEX14-MT Interface. (A) NMR structure (PDB: 2W84) of PEX14-NTD (purple) in complex with a PEX5-derived peptide (blue). The sidechains of PEX14 residues F52 and K56 at the PEX5-binding interface and K54 that is not involved in PEX5 binding are shown. (B, C) Analysis of MT binding of PEX14 variants harboring mutations in the putative binding region. GST-PEX14-NTD variants (F52A, K54A, K56A, and F52A/K56A) were purified and tested in ELISA plate assays with coated MT (B, C). F52 as well as K56 were identified as critical residues for the interaction with PEX5 and MT whereas the K54A mutation did not affect the binding affinities of PEX14-NTD, consistent with the binding interface also seen with PEX5. The experimental data were fitted with standard one-site models using non-linear regression and shown with standard errors (n=2).

### The C-terminal region of ß-tubulin contains two PEX14-binding motifs

To identify the PEX14-NTD-interacting site of ß-tubulin, purified GST-PEX14-NTD was incubated with a membrane-bound peptide library representing the 444 amino acid sequence of human tubulin ß5, the isoform with the highest score in the mass-spectrometric analyses of affinity-purified PEX14 complexes [12]. To this end, 15-mer peptides with a 13-amino acid overlapping region were synthesized on membrane supports. Immunodetection of bound GST-PEX14-NTD revealed two binding regions: a region involving amino acids 187–211 and a second site close to the ß-tubulin C-terminus involving amino acids 391–431 located on helices H11 and H12 (**Fig. 4A**). Interestingly, in the three-dimensional structure these two regions are in close proximity (**Fig. 4B**). The region comprising amino acids 187–211 forms hydrophobic contacts with the C-terminal helical hairpin [27]. The antiparallel hairpin build by the two C-terminal helices together with the glutamate-rich C-terminal end, referred to as E-hook, are located on the exterior of the microtubule and together with parts of α-tubulin form the contact zone for interaction with the motor proteins kinesin and dynein [28]. Since the PEX14-interacting peptides of ß-tubulin located within the N-terminal binding region are only partially accessible and show weaker affinity to PEX14-NTD, we focused on PEX14 interaction with the C-terminal region of ß-tubulin.

**Figure 4.**
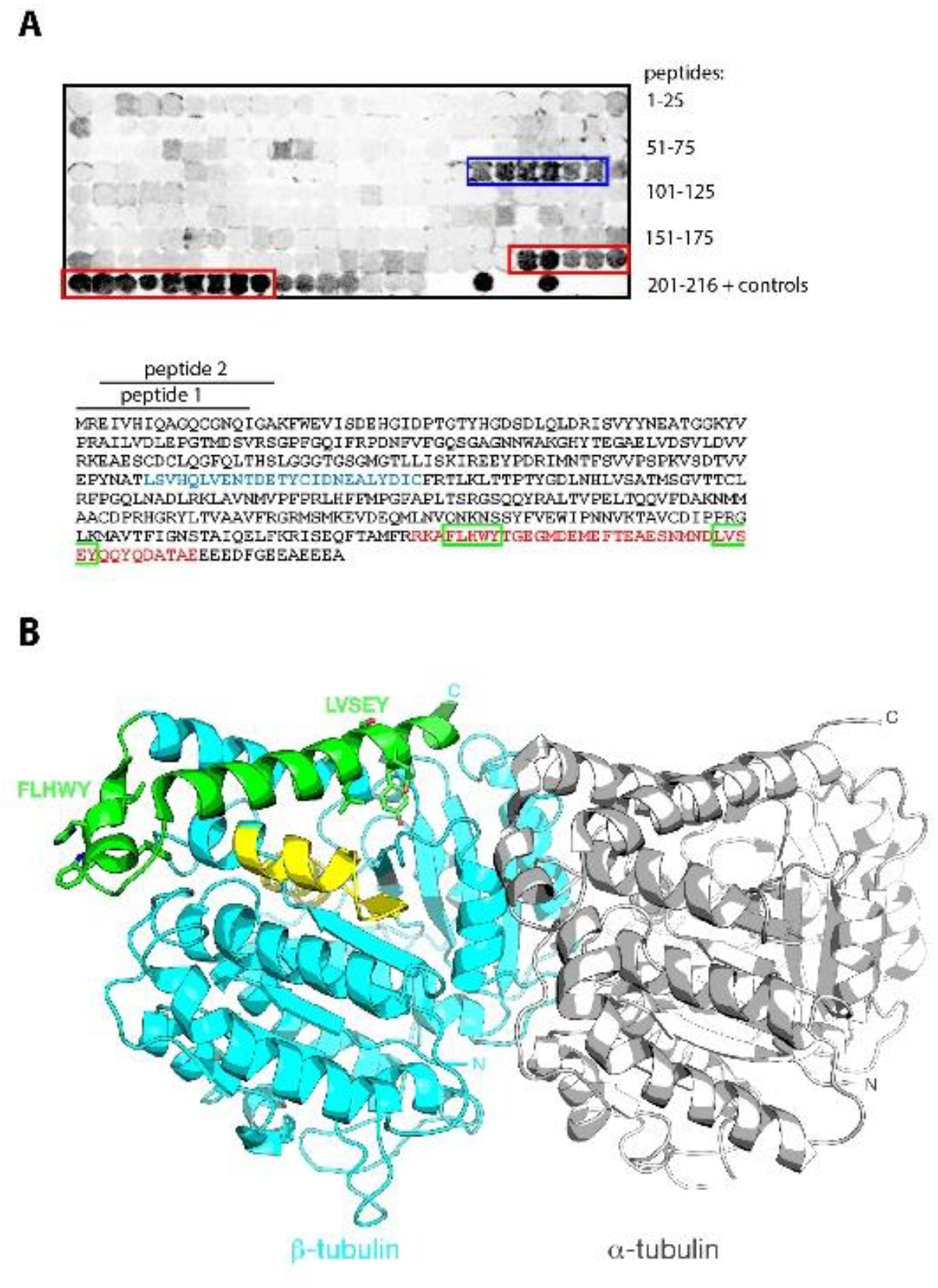
Identification of the PEX14-NTD-binding site in ß-tubulin. (A) Full-length sequence of human ß-tubulin (tubulin beta-5, UniProt P07437, 444 amino acids) was synthesized as 15-mer peptides with an overlap of 13 amino acids. Each spot on the membrane represents one immobilized peptide beginning with the N-terminal 15 amino acids (peptide 1). As controls, two additional 15-mer peptides representing human PEX5 (aa 57–71 and aa 113–127) were applied, each containing a high-affinity PEX14-binding pentapeptide motif, either LVAEF or WxxxF/Y sequence. After incubation with purified GST-PEX14-NTD, interacting peptide spots were detected with anti-GST antibodies. Interacting tubulin peptides are marked by blue and red boxes and the corresponding amino acid sequences (aa 187–211 and aa 391–431) are highlighted in the sequence shown below. The green boxes within the C-terminal sequence region highlight two WxxxF/Y-like sequence motifs. (B) The structure of α-tubulin (gray) and ß-tubulin (cyan) (PDB: 1JFF). The PEX14-NTD binding regions corresponding to aa 187–211 and 391–431 of human ß-tubulin are colored in yellow and green, respectively. Sidechains of two PEX14-binding motifs in the C-terminus of ß-tubulin are shown.

In the C-terminal region of ß-tubulin two putative PEX14-binding motifs are of interest. These are the pentapeptides FLHWY (aa 394–398) and LVSEY (aa 418–422), of which the latter is very similar to the first PEX14-binding motif in human PEX5 with the sequence LVAEF [24]. Consistently, a protein fragment corresponding to the C-terminal region of ß-tubulin (aa 388–444) binds to two molecules of PEX14-NTD as shown by isothermal titration calorimetry (ITC) (**Supplementary Figure 3**).

To investigate the interaction of PEX14 with ß-tubulin at the residue-specific level, NMR titration experiments were performed using ^15^N-labeled PEX14-NTD and two unlabeled 15-mer tubulin peptides that harbor the pentapeptide binding motifs FLHWY and LVSEY. In both cases, clear chemical shift perturbations were observed upon addition of the peptide to PEX14-NTD, indicating that each motif can interact with PEX14-NTD (**Fig. 5A, B**). Binding of the LVSEY peptide is mostly in the fast-exchange regime on the NMR chemical shift timescale as judged from the gradual shift in NMR peak positions with increasing peptide concentrations. Few residues, such as L58, exhibit intermediate-exchange binding as the peaks disappear and reappear at higher peptide concentrations. In the case of the FLHWY peptide, however, binding is exclusively in the fast-exchange regime and fewer residues experience chemical shift perturbations, indicative of a weaker interaction (**Fig. 5**). Based on the chemical shift perturbations, apparent *K_D_* values of ~5.0 and ~282 μM were estimated for the LVSEY and FLHWY peptides, respectively (**Fig. 5C, D**). These values are at least 50-fold weaker than the *K_D_* of ~100 nM reported for the interaction of PEX14-NTD with the first WxxxF/Y motif in human PEX5 [23], providing a rationale for the competition of PEX5 with MT for binding to PEX14.

**Figure 5.**
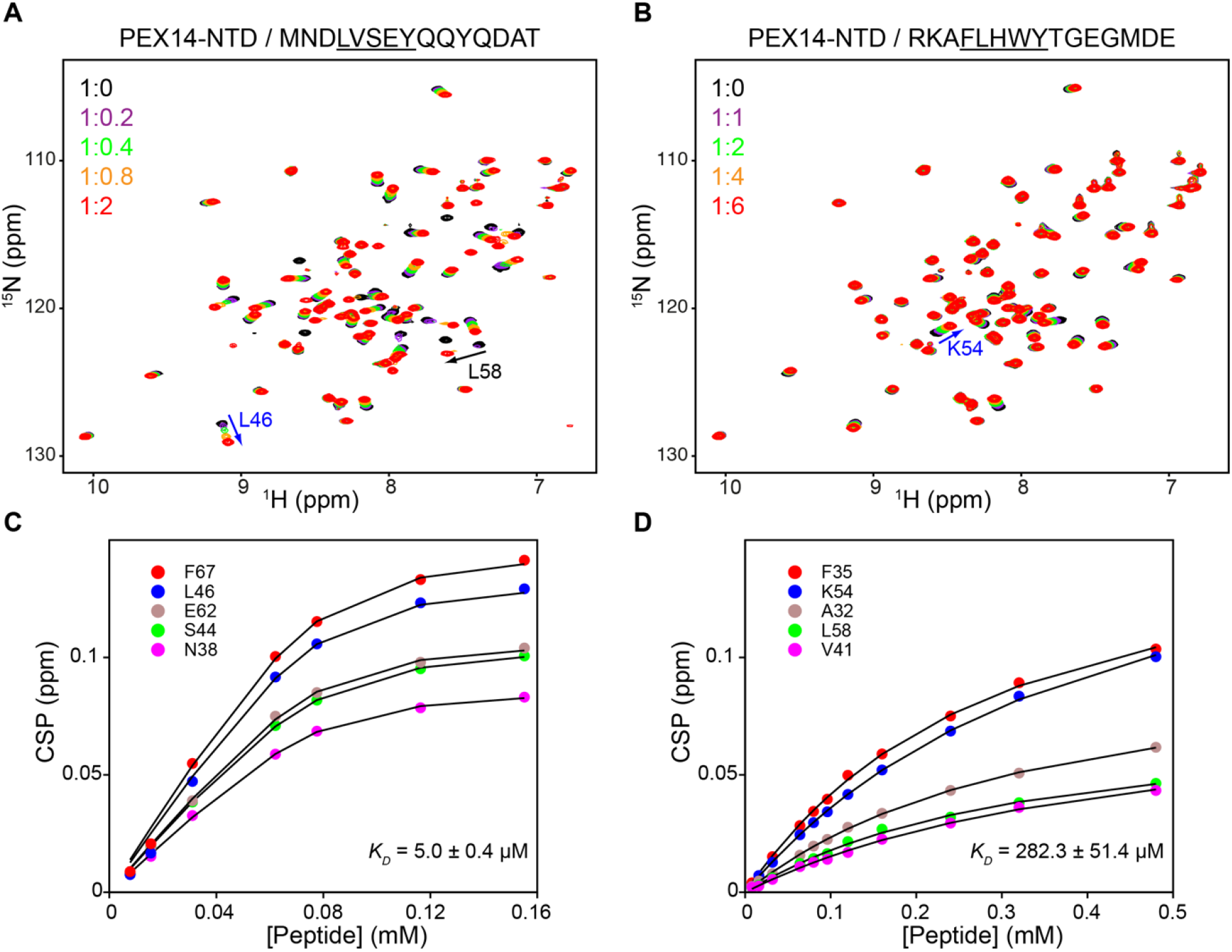
NMR analysis of the PEX14-tubulin interaction. NMR titration of PEX14-NTD with (A) LVSEY and (B) FLHWY peptides derived from the human tubulin β-chain. Overlay of ^1^H–^15^N HSQC spectra of free ^15^N-labelled PEX14-NTD (black) and in the presence of increasing concentrations of tubulin peptide (magenta, green, orange and red). Selected residues undergoing chemical shift perturbation are indicated. Five residues undergoing fast-exchange binding to (C) LVSEY and (D) FLHWY peptides were selected and the corresponding binding curves were fitted to obtain the *K_D_*.

To further analyze the structural basis of the competitive effects of PEX5 on PEX14-tubulin interactions, an NMR double-titration experiment was performed in which the unlabeled LVSEY peptide was added to saturation to ^15^N-labeled PEX14-NTD followed by addition of a PEX5 peptide corresponding to the first WxxxF/Y motif with the sequence ALSEN<u>WAQEF</u>LAAGDA (referred to as the WAQEF peptide). Addition of only 0.5 molar equivalent of the WAQEF peptide to the PEX14-LVSEY complex resulted in drop in intensity of peaks that had shifted upon LVSEY binding and appearance of new signals corresponding to the PEX14-WAQEF complex (**Fig. 6A**). This is indicative of slow exchange binding of the WAQEF peptide, consistent with the nanomolar *K_D_* value. It also demonstrates that both the tubulin-derived LVSEY and the PEX5-derived WAQEF bind to the same binding site on PEX14. Further addition of the WAQEF peptide completely shifted the spectrum of PEX14-NTD to the WAQEF-bound form, indicating that the WAQEF peptide can replace the LVSEY peptide (**Fig. 6B**).

**Figure 6.**
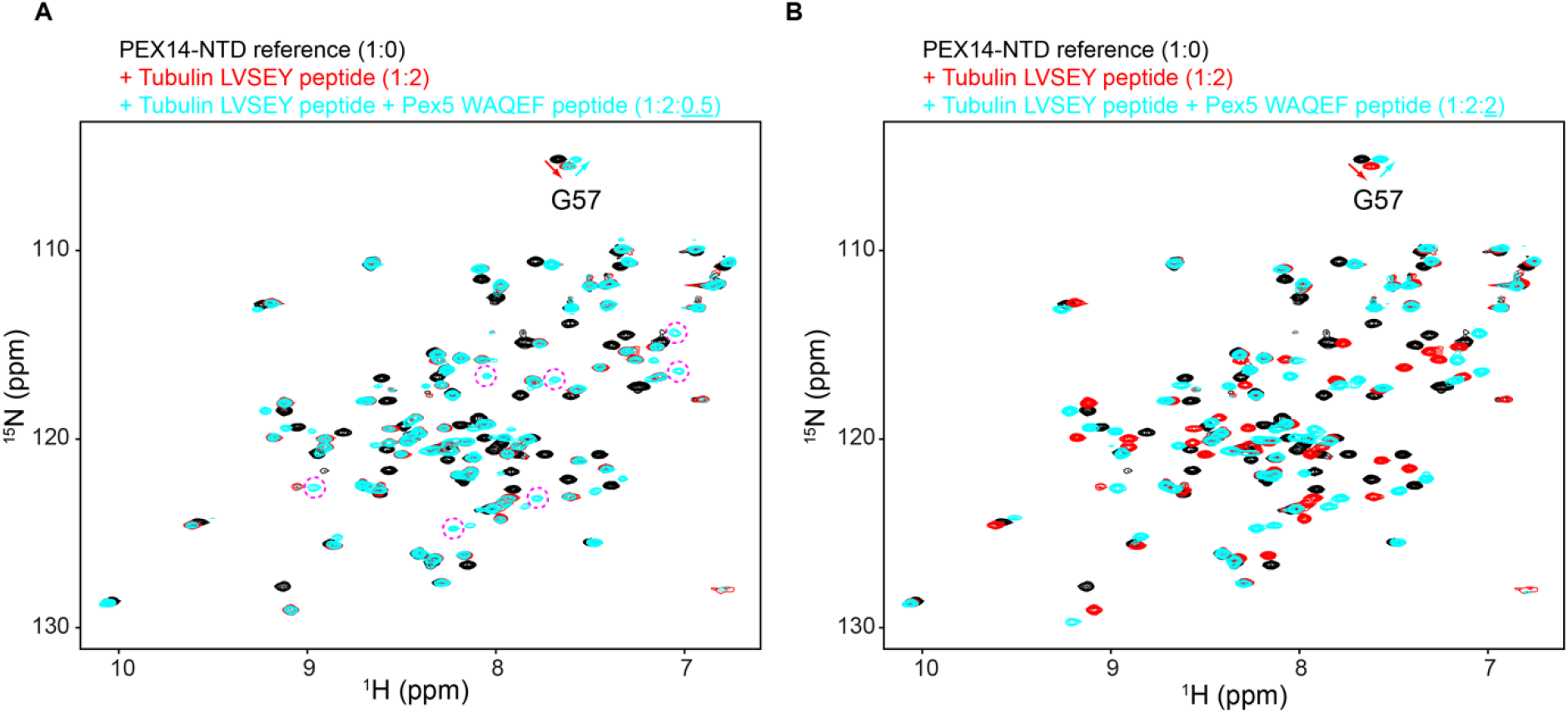
NMR competition experiments. ^1^H–^15^N HSQC spectra of ^15^N-labelled PEX14-NTD free (black), in the presence of two-fold molar excess of the LVSEY peptide (red), and in the presence of both LVSEY and WAQEF peptides (cyan). (A) Addition of 0.5 molar equivalent of the WAQEF peptide to the PEX14-LVSEY complex (1:2 stoichiometry) results in appearance of new signals (selected peaks are encircled) corresponding to the PEX14-WAQEF complex. (B) At equal concentrations, the WAQEF peptide completely displaces the LVSEY peptide, thus shifting the spectrum of PEX14-NTD to the WAQEF-bound form. The peak corresponding to G57 is annotated. LVSEY and WAQEF refer to peptides derived from the tubulin β-chain and PEX5, respectively.

Taken together, these data indicate that human PEX14-NTD interacts with a region comprising two binding-motifs in the C-terminus of tubulin β-chain. Interestingly, one of the motifs (LVSEY) is highly similar to the LVxEF motif in PEX5 and is therefore expected to have the same mode of binding to PEX14.

### PEX14-bound MT show reduced affinity to the kinesin motor domain

Since both PEX14 and the kinesin motor domains bind to the C-terminal domain of ß-tubulin in close proximity, we asked whether they compete for binding. One possible scenario is that binding of motor proteins releases the potential peroxisome-tethering PEX14 to allow movement along MT. To test this hypothesis, we applied purified kinesin motor domain to the MT-PEX14 *in vitro* binding assay. The recombinant kinesin fragment is a monomeric variant of KIF5B, a member of the kinesin 1 family that has previously been used in binding assays [29]. The His-tagged motor domain of KIF5B, which is well conserved in the kinesin superfamily, contains the MT-binding site and an ATP-hydrolysis site. We used the plate-ELISA assay to estimate the affinity of this fragment of KIF5B to MT. The *K_D_* values for the interaction are in the nanomolar range (56.6 +/- 15.6 nM, **Fig. 7A, B**), but significantly higher than that of GST-PEX14 (*K_D_* of 0.7–2.3 nM, **Fig. 1A, Fig. 7C**). The binding constants for the kinesin–MT interaction are in good agreement with published values for other monomeric kinesin 1 variants [30–32].

**Figure 7.**
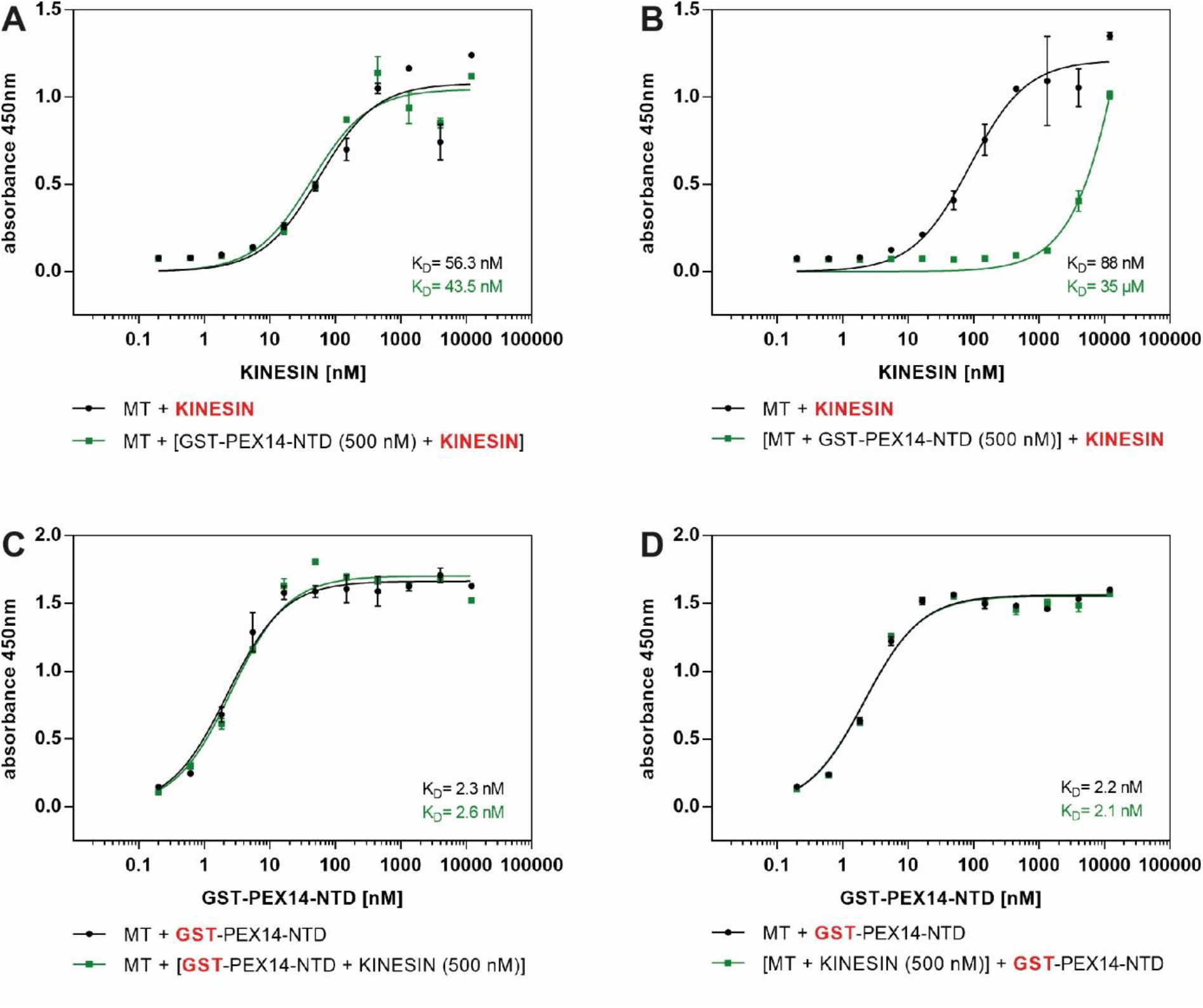
Analysis of the correlation of kinesin and PEX14-binding to MT. Bound MT were incubated with various concentrations of His-tagged motor protein domains of monomeric KIF5B (KINESIN) or GST-PEX14-NTD for 1 h at room temperature as indicated. Binding of kinesin to MT in the simultaneous presence of 500 nM GST-PEX14-NTD (A) or binding of kinesin to MT that were pre-incubated with 500 nM GST-PEX14-NTD (B) was monitored. Binding of GST-PEX14-NTD to MT in the simultaneous presence of 500 nM kinesin (C) or binding of GST-PEX14-NTD to MT that were pre-incubated with 500 nM kinesin (D) was recorded. Primary antibodies used for immunodetection of bound kinesin and PEX14 were anti-His and anti-GST, respectively. The experimental data were fitted with standard one-site models using non-linear regression and shown with standard errors (n=2).

To test possible effects of PEX14 on kinesin binding to MT, we analysed the binding of the kinesin motor domain in the presence of GST-PEX14-NTD. While the simultaneous presence of 500 nM PEX14 in the binding assay did not affect the association of kinesin with MT (**Fig. 7A**), one hour preincubation of MT with 500 nM GST-PEX14-NTD drastically reduced the MT-binding affinity of kinesin by at least three orders of magnitude (**Fig. 7B**). These data suggest that pre-bound GST-PEX14-NTD impairs MT-binding of the kinesin motor domain. Residual binding of kinesin under these conditions might be explained by the large interface formed between the kinesin motor domain and both α- and ß-tubulin subunits [28, 30, 32]. PEX14 competition at the C-terminal part of the ß-subunit might not interfere with binding of the motor domain to the α-tubulin subunit.

On the other hand, neither the simultaneous presence of 500 nM kinesin (**Fig. 7C**) nor saturation of MT with 500 nM kinesin (**Fig. 7D**) affected MT-binding of GST-PEX14-NTD. These results argue against a simple dislocation mechanism in which the motor domain can replace the potential tethering factor PEX14 by direct competition. Collectively, our results suggest that under physiological conditions PEX14-NTD and the kinesin motor domain can bind to MT simultaneously.

## Discussion

The peroxisomal membrane protein PEX14 has been proposed as an initial anchor point for the association of the organelles at MT [4, 12]. Here, we studied the structural basis of interaction of the human PEX14 and microtubular filaments in molecular detail. We found that the N-terminal domain of PEX14, which is also the docking site for peroxisomal import receptors, interacts with MT. Interestingly, the molecular requirements for the MT-binding of PEX14 are essentially identical for its interaction with PEX5, the cytosolic import receptor for peroxisomal matrix enzymes. On the PEX14 side, we showed that mutation of residues that are critical for PEX5 binding also reduces the affinity for MT. On the MT side, we identified two PEX14-interacting alpha helical pentapeptide motifs within the C-terminal region of the ß-subunit of tubulin, similar to those that mediate interactions between peroxisomal import receptors and PEX14. Notably, single PEX14-binding peptides of tubulin exhibit much lower affinity when compared with single PEX5-derived peptides and consequently, full-length PEX5, with eight PEX14-binding motifs, is a strong competitor of PEX14-MT interaction both *in vitro* and *in vivo*.

What might be the functional meaning of the PEX5 competition on MT-binding of PEX14 in a cellular environment? It has been shown that PEX14 is indispensable for Miro1 splicing variant 4 -dependent motility of peroxisomes without directly interacting with the motor protein adaptor complex [10]. However, PEX14 tethering to MT might be a precondition to enable the organization of motor proteins and their adapters at the organellar surface [12]. Our data suggest that PEX5 blocks the ß-tubulin interaction site of PEX14 and thereby competes peroxisome attachment at MT. Consistent with this, overexpression of the peroxisomal import receptor PEX5 in human cells interferes with the motility of peroxisomes (**Fig. 2**). Such a correlation of peroxisomal protein import and microtubular -activities could enable directional placement of peroxisomes. Moreover, it has been suggested that cargo-loading significantly increases docking of PEX5 at the peroxisomal membrane [33–35]. For the functional relevance of our results, this means that high abundance of cargo-loaded import receptors that associate with PEX14 block its binding site for ß-tubulin and thus interfere with its association with MT thereby preventing peroxisomal movements. A lowering of the local concentration of cargo would deplete the receptor from the membrane and release the PEX14 tubulin binding site, which is now available for binding to MT. Subsequent binding of motor proteins is supposed to move peroxisomes to cellular areas with higher concentration of receptor-cargo complexes, maybe regions with a lower abundance of import-active peroxisomes. This scenario allows the regulated cellular distribution of peroxisomes. This model is in-line with previous observations by Nguyen et al showing that peroxisomal remnants in many PEX1-deficient Zellweger cell lines exhibit clustering and loss of alignment along MT [36]. The lack of PEX1, which is required for recycling of the import receptor from the membrane, results in an accumulation of PEX5 and occupancy of PEX14 at the peroxisomal membrane [37]. As a result and in accordance with the model, microtubular tethering and peroxisomal motility would be blocked.

Movement of peroxisomes along MT requires binding of motor proteins and dissociation of the tethering complex. Motor proteins like dynein or kinesin as well as adaptor proteins like dynactin bind to the C-terminal region of ß-tubulin, especially to the flexible C-terminal E-hook [28, 38, 39]. Notably, one of the PEX14-interacting peptides that we have identified here overlaps with the kinesin binding site of ß-tubulin [30]. Importantly, substitution of the E within the LVSEY motif was shown to significantly reduce the affinity of MT for kinesin [30]. Therefore, we asked whether a simple displacement strategy based on competition of PEX14 and kinesin underlies dissociation of the peroxisomal MT-tethering complex. Consistent with the close proximity of the binding sites, we found that GST-PEX14-NTD hinders MT-binding of kinesin at high concentrations. However, the presence of kinesin, even at MT-saturating concentrations showed no direct effect on PEX14 association with MT. This suggests that the binding of motor proteins alone is not sufficient to dissociate the PEX14 MT-tethering complex. In a cellular environment, manifold posttranslational modifications are involved in the regulation of organelle association with MT [40]. These remain to be identified in the context of peroxisome motility.

## Material and Methods

### Plasmids

The construction of the bicistronic expression vector coding for both *Hs*PEX5 and EGFP-*Hs*PEX26 was based on the previously described *Hs*GFP–PEX26 expression plasmid [41] and a bicistronic expression vector coding for *Hs*PEX5 as well as eGFP-SKL [24]. The DNA-sequence encoding *Hs*PEX26 was PCR-amplified and subcloned into pIRES-eGFP digested by NotI and BsrGI.

*E.coli* expression plasmids encoding GST-PEX14-NTD with amino acid substitutions were constructed via Site Directed Mutagenesis using QuikChange II (Stratagene). ß-tubulin (amino acids 388–444) was cloned into the pProEx HTa expression vector (Life Technologies). All oligonucleotides used are listed in Supplementary Table 1.

*E. coli* GST-PEX14-NTD (GST-MTBD) expression plasmid [22] and full-length His_6_-TEV-PEX5S [42] have been described previously.

*E. coli* RP hk339-GFP (monomeric His-tagged Kif5B kinesin motor domain) expression plasmid was a gift from Ron Vale (Addgene plasmid #24431).

### Expression and purification of recombinant proteins

Expression of full length His_6_-PEX5, PEX14-NTD, ß-tubulin(388–444) and His_6_-tagged kinesin motor domain was carried out in *E. coli* strain BL21(DE3). The proteins were expressed in LB media supplemented with 50 mg/l kanamycin or 100 mg/l ampicillin. After incubation at 37 °C, cells were induced in mid-log phase with 0.4 mM IPTG for PEX5 and PEX14 and with 0.1 mM IPTG for kinesin. Cells were then further cultivated for 4 hours for PEX5 and PEX14 and overnight for the kinesin constructs. Cells were harvested and stored at −80 °C. Cells were disrupted by Emulsiflex in 50 mM Tris-HCl, pH 7.4, 150 mM NaCl and 1 mM DTT for PEX5, or in 50 mM Tris-HCl, pH 8.0, 150 mM NaCl and 1 mM DTT for PEX14-NTD forms, or in 250 mM NaCl, 1 mM MgCl2, 0.5 mM ATP, 50 mM phosphate buffer, pH 8.0 for kinesin, plus protease inhibitors (8 mM Antipain, 0.3 mM Aprotinin, 1 mM Benzamidine hydrochloride, 1 mM Bestatin, 10 mM Chymostatin, 5 mM Leupeptin, 5 mM Sodium Fluoride, 15 mM Pepstatin A, 1 mM PMSF). After sedimentation of the cells debris at 3000 x g for 20 min, the supernatants were collected as cell lysates and incubated with Ni-NTA agarose or glutathione-Sepharose for 1 hour at 4 °C. Bound proteins were transferred to a column and washed with 200 ml of washing buffer (lysis buffer without inhibitors). His_6_-tagged proteins were eluted with 50 mM to 500 mM imidazole and GST-tagged proteins with 1 mM to 10 mM glutathione in 1 ml fractions. All affinity purification steps of the different proteins were monitored by SDS-PAGE analysis.

For further purification, PEX5 was applied to mono Q ion-exchange chromatography using Äkta Purifier HPLC system (GE Healthcare). The column was equilibrated with 500 mM NaCl, pH 8.0, 1 mM DTT. Bound proteins were eluted with a linear gradient of 0 mM to 500 mM NaCl, collected in 0.5 ml fractions and analyzed by SDS-PAGE and Coomassie staining.

Purification of porcine brain tubulin was carried out as described previously [43].

### Peptide spot overlay assays

Human ß-tubulin (Tubulin beta-5) were synthesized as peptide libraries comprising peptides with a length of 15 amino acids and overlapping regions of 13 amino acids. Peptides were directly synthesized on a cellulose membrane as described previously [25, 44]. After blocking with 3% BSA in TBS (10 mM Tris/HCl pH 7.4, 150 mM NaCl), membranes with immobilized tubulin peptides were probed overnight at 4°C with purified 10 nM GST-PEX14-NTD in TBS. Bound PEX14-NTD was immunodetected with monoclonal anti-GST antibodies (Sigma-Aldrich) in TBS plus 3% BSA, and fluorescent-labelled secondary antibodies in TBS plus 10% milk powder by Licor Odyssey scanning. Between these steps, the membranes were thoroughly washed with TBS-TT (20 mM Tris/HCl pH 7.5, 0.5 M NaCl, 0.05% (v/v) Tween20, 0.2% (v/v) Triton X-100) and at the end of each washing step with TBS only.

### In vitro binding assay

10 μg of either purified porcine brain tubulin or PEX5 in phosphate-buffered saline (PBS) were bound to 96-well immunoplates (Immulon™ 2 HB) for 1 hour. After 1 hour blocking with 3% BSA (250 μl), wells were washed two times with 280 μl PBS (pH 7.5). As analytes, GST- or His-tagged PEX14-NTD, kinesin or PEX5 in PBS were added to the coated protein in dilution series ranging from 3 μM to 0.05 nM or with a fixed concentration of 500 nM to the coated protein, followed by three washing steps with PBS + 0.05% Tween20 (pH 7.5). For analysing competitive effects on the PEX14-MT interaction, increasing concentrations of PEX14-NTD were pre-incubated with coated MT before adding 500 nM of PEX5 or kinesin to the plate. Alternatively, both proteins together were incubated with immobilized MT for 1 hour at RT, followed by three washing steps with PBS+0.05% Tween. For the immunological detection of bound proteins, 100 μl of monoclonal mouse-anti-GST antibodies (Sigma-Aldrich), monoclonal anti-His_6_ antibodies (ThermoFisher) or polyclonal anti-PEX14 antibodies [45] in PBS+0.05% Tween were used. After 1 hour incubation at RT the plates were washed three times with PBS+0.05% Tween each, followed by incubation with secondary antibodies such as horse radish peroxidase (HRP)-conjugated anti-mouse antibody (ThermoFisher Scientific) or HRP-conjugated anti-rabbit antibody (Dako), followed again by three washing steps with PBS+0.05% Tween. TMB (Tetramethylbenzidine) and hydrogen peroxidase (1:1) were added to the interacting proteins. After 3 to 10 min the color reaction was halted by adding 1N sulfuric acid and absorbance was measured at a wavelength of 450 nm.

### Live cell imaging

For live cell imaging, cell lines were transfected with bicistronic plasmids encoding eGFP-*Hs*PEX26 or PEX5/eGFP-*Hs*PEX26 and cultivated for 48 hours as described previously [46]. Overexpression of PEX5 was controlled by immunoblot analysis of cell lysates using anti PEX5 antibodies [47] and anti-Prohibitin antibodies (Abcam) as loading control. Confocal fluorescence microscopy was performed using the Zeiss LSM510 Meta (Zeiss, Oberkochen) with the oil immersion objective Plan-Neofluar 40x/1.3 (Zeiss, Oberkochen). The CTI Controller 3700 regulated the air temperature with its associated tempcontrol minisystem (PeCon, Erbach). For time-lapse imaging, the argon ion laser of the LSM510 Meta stimulated the fluorescent proteins at a wavelength of 488 nm. The cells were imaged at a threefold magnification and with a resolution of 512 x 512 Pixels. Finally, the processing and analysis of gained images was conducted with the Zeiss LSM Image Browser Version 4.2.0.121 [48]. The effect of overexpression of *Hs*PEX5 on the motility of peroxisomes was quantitatively analyzed by seven independent experiments. SPSS Statistics and GraphPad Prism 8 software were used for statistical analysis. Due to lack of homogeneity of variance, a Mann-Whitney-U-Test was performed to test the null hypothesis. Error bars show the standard error. P values lower than 5% were determined as significant differences.

### ITC

ITC titrations were performed on an iTC200 microcalorimeter at 25 °C. PEX14-NTD at 438 μM concentration in the syringe was injected into the cell containing 52 μM ß-tubulin(388–444). The ITC thermogram was integrated using NITPIC [49] and the data were fitted to a non-symmetric two binding-sites model in SEDPHAT [50].

### NMR spectroscopy

NMR experiments were recorded at 298 K on a 600-MHz Bruker spectrometer equipped with a cryogenic probe. Spectra were processed using NMRPipe [51] and analyzed using SPARKY 3 (T. D. Goddard and D. G. Kneller, University of California, San Francisco). NMR CSPs as a function of total peptide (ligand) concentration were fitted to the following equation to derive equilibrium dissociation constants: 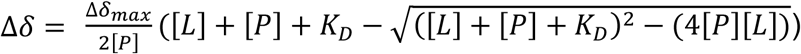 where Δδ is the combined ^1^H and ^15^N CSP at a given ligand concentration [*L*], and [*P*] is the protein concentration. The *K_D_* values reported are averages over five residues plus standard deviation.

### NMR sample preparation

^15^N-PEX14(16–80W) was expressed and purified as described previously [24]. The NMR buffer consisted of 20 mM sodium phosphate at pH 6.5, 100 mM NaCl and 1 mM DTT. Peptides of human PEX5 (ALSENWAQEFLAAGDA) and ß-tubulin (RKAFLHWYTGEGMDE; MNDLVSEYQQYQDAT) were chemically synthesized (Peptide Specialty Laboratories GmbH) with >95% purity. NMR titration were done by adding increasing amounts of peptide solution to ^15^N-PEX14 at 80 μM concentration.

## Supplements

**Supplementary Table 1.**
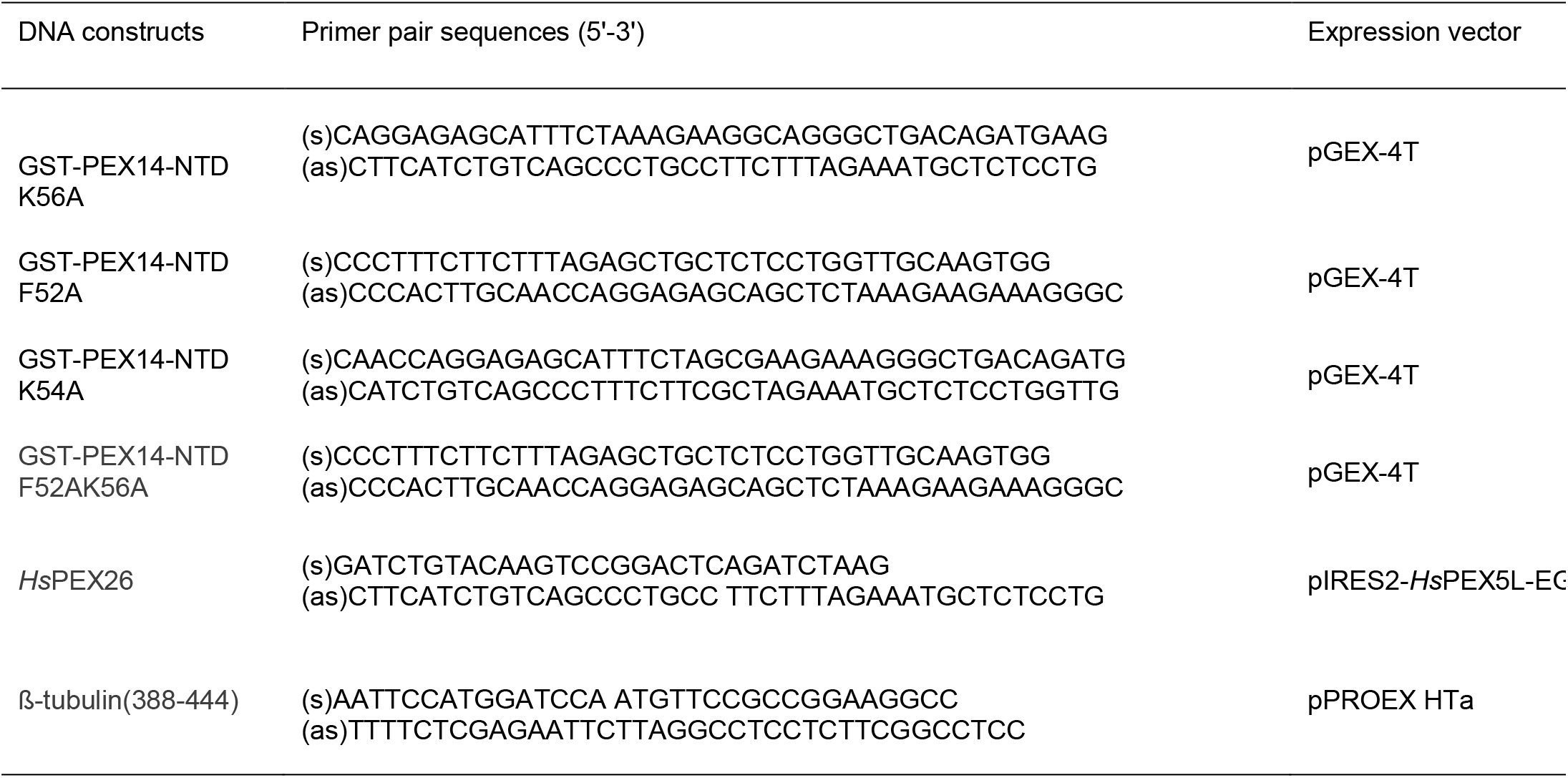
Oligonucleotides used in this study. Oligonucleotides (s), sense; (as), antisense

**Supplementary Figure 1.**
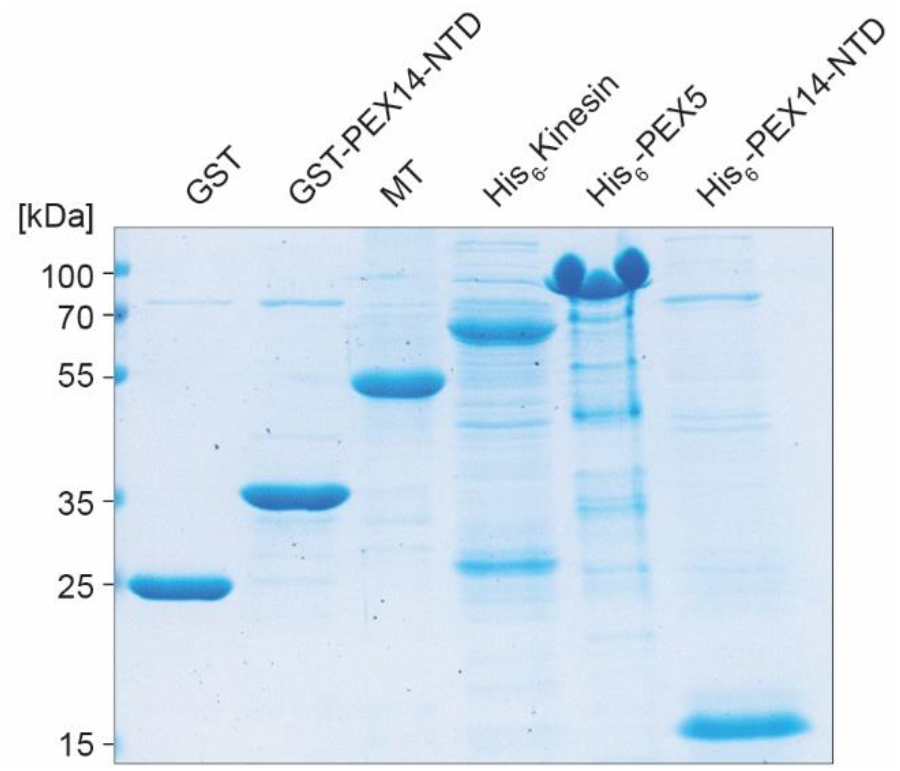
Purification of proteins used in the ELISA MT-binding assays. GST, GST-PEX14-NTD, His_6_-tagged kinesin, PEX5 and PEX14-NTD were expressed in *E.coli* and affinity-purified. MT were isolated from porcine brain homogenates using polymerizing/depolymerizing cycles and centrifugation steps. Enriched protein fractions containing 5 nmol protein each were analyzed by SDS-PAGE and Coomassie staining.

**Supplementary Figure 2.**
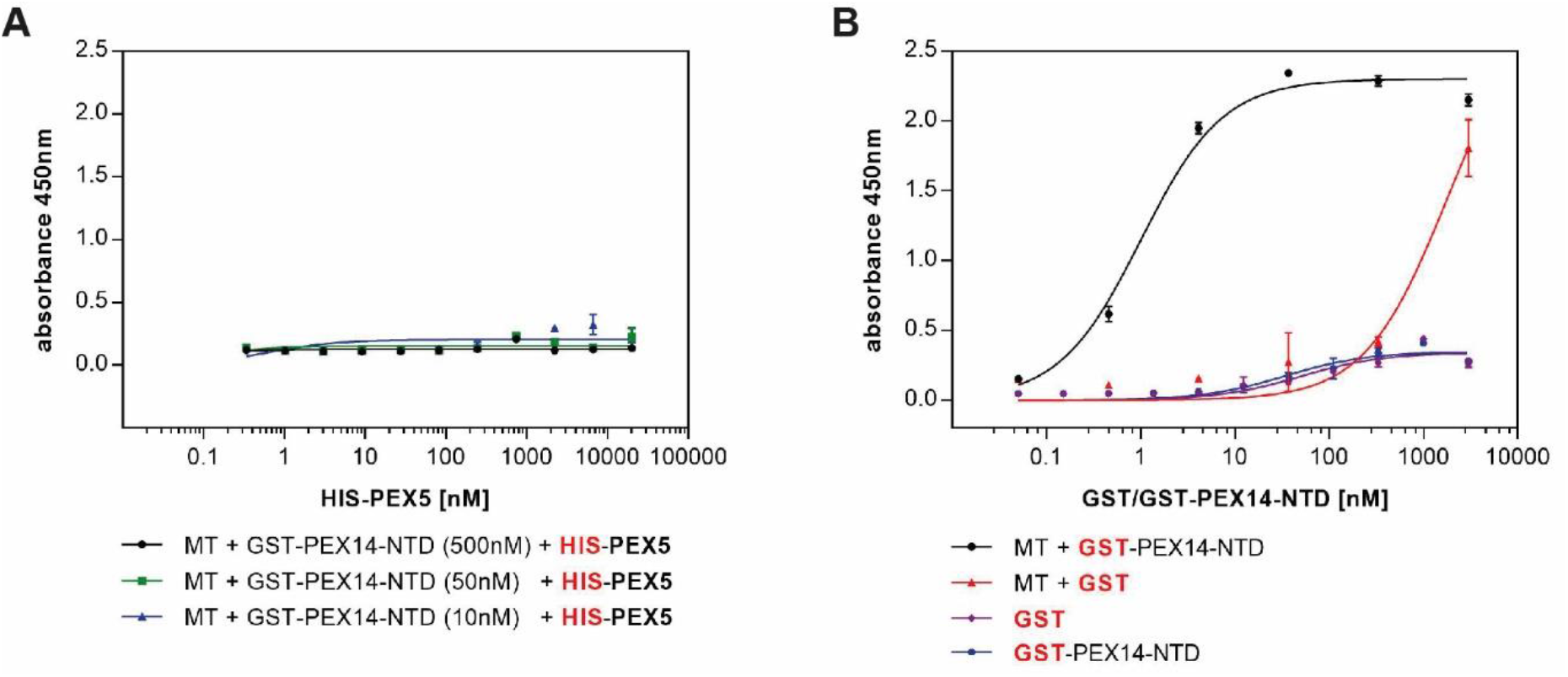
Control experiments for the ELISA MT-binding assay. (A) Microtiter plates were coated with MT and incubated with three different concentrations of GST-tagged PEX14-NTD (10, 50 and 500 nM) and varying concentrations of His-tagged PEX5. Neither His-tagged PEX5 nor PEX5/GST-PEX14-NTD complex interact with immobilized MT. (B) Microtiter plate wells which were coated with MT and milk powder as blocking reagent or incubated with blocking solution alone were tested with varying concentrations of GST-PEX14 and GST alone. GST alone showed unspecific binding to MT at protein concentrations in the μM range. Bound proteins were immunodetected by specific antibodies against His-tag (A) or GST (B). The experimental data were fitted to a standard one-site model using non-linear regression and shown with standard errors (n=2).

**Supplementary Figure 3.**
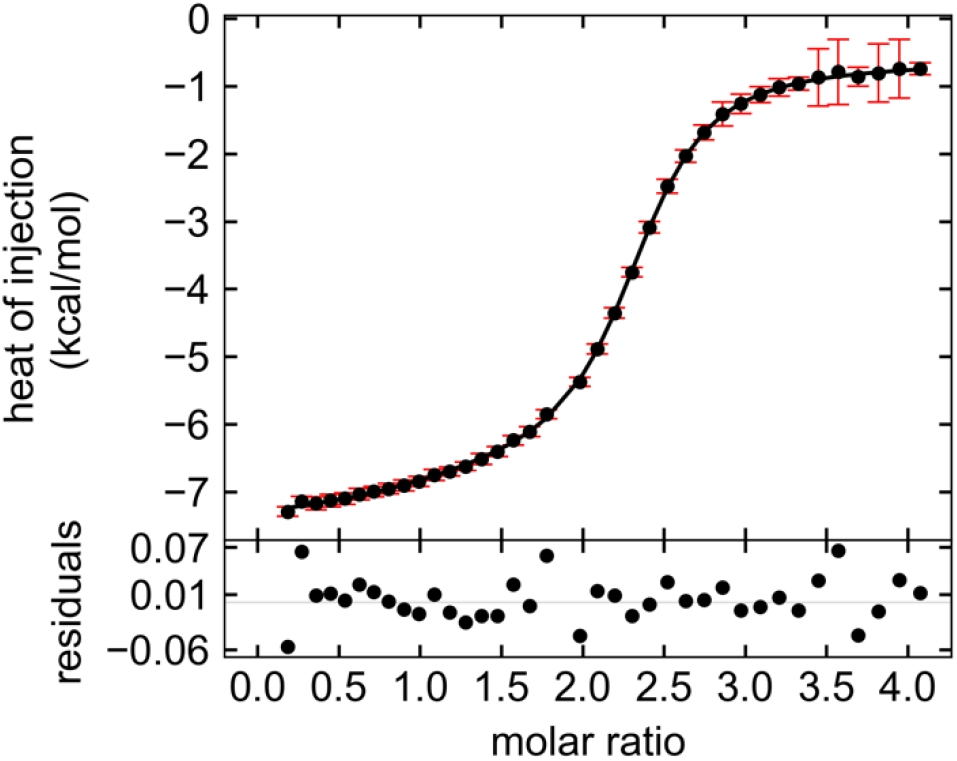
ITC data of ß-tubulin(388–444) and PEX14-NTD interaction. PEX14-NTD in the syringe was injected into the ITC cell containing recombinant ß-tubulin(388–444). The data were best fitted to a non-symmetric two binding-sites model. The bottom panel shows the residuals of the fit. The 1:2 stoichiometry of the interaction is also evident from the molar ratio that corresponds to the mid-point of the binding curve.

## Acknowledgements

This work was supported by the Deutsche Forschungsgemeinschaft (FOR1905) to R.E. and M.S.

## Author contributions (CRediT author statement)

**Maren Reuter**: Conceptualization, Methodology, Investigation, Formal analysis, visualization, Writing and Editing

**Hamed Kooshapur**: Conceptualization, Methodology, Investigation, Formal analysis, Visualization, Writing and Editing

**Jeff-Gordian Suda**: Investigation and Formal analysis, Visualization

**Alexander Neuhaus**: Methodology, Investigation, Formal analysis

**Lena Brühl**: Investigation and Formal analysis

**Pratima Bharti**: Investigation and Formal analysis

**Martin Jung**: Resources

**Wolfgang Schliebs**: Conceptualization, Supervision, Visualization, Writing and Editing

**Michael Sattler:** Conceptualization, Writing and Editing, Project administration and Funding acquisition

**Ralf Erdmann**: Conceptualization, Supervision, Writing and Editing, Project administration and Funding acquisition

## Abbreviations

GST: glutathione S-transferase
MT: microtubules
NTD: N-terminal domain
PBD: peroxisome biogenesis disorder
PEX: peroxins
TEV: tobacco etch virus protease cleavage site

## References

[1] Islinger M, Voelkl A, Fahimi HD, Schrader M. The peroxisome: an update on mysteries 2.0. Histochem Cell Biol. 2018;150:443–71.

[2] Schrader M, Grille S, Fahimi HD, Islinger M. Peroxisome interactions and cross-talk with other subcellular compartments in animal cells. Subcell Biochem. 2013;69:1–22.

[3] Sugiura A, Mattie S, Prudent J, McBride HM. Newly born peroxisomes are a hybrid of mitochondrial and ER-derived pre-peroxisomes. Nature. 2017;542:251–4.

[4] Neuhaus A, Eggeling C, Erdmann R, Schliebs W. Why do peroxisomes associate with the cytoskeleton? Biochimica et biophysica acta. 2016;1863:1019–26.

[5] Rapp S, Saffrich R, Anton M, Jakle U, Ansorge W, Gorgas K, et al. Microtubule-based peroxisome movement. J Cell Sci. 1996;109 (Pt 4):837–49.

[6] Wiemer EA, Wenzel T, Deerinck TJ, Ellisman MH, Subramani S. Visualization of the peroxisomal compartment in living mammalian cells: dynamic behavior and association with microtubules. J Cell Biol. 1997;136:71–80.

[7] Kural C, Kim H, Syed S, Goshima G, Gelfand VI, Selvin PR. Kinesin and dynein move a peroxisome in vivo: a tug-of-war or coordinated movement? Science. 2005;308:1469–72.

[8] Castro IG, Schrader M. Miro1 - the missing link to peroxisome motility. Commun Integr Biol. 2018;11:e1526573.

[9] Castro IG, Richards DM, Metz J, Costello JL, Passmore JB, Schrader TA, et al. A role for Mitochondrial Rho GTPase 1 (MIRO1) in motility and membrane dynamics of peroxisomes. Traffic. 2018;19:229–42.

[10] Okumoto K, Ono T, Toyama R, Shimomura A, Nagata A, Fujiki Y. New splicing variants of mitochondrial Rho GTPase-1 (Miro1) transport peroxisomes. J Cell Biol. 2018;217:619–33.

[11] Dietrich D, Seiler F, Essmann F, Dodt G. Identification of the kinesin KifC3 as a new player for positioning of peroxisomes and other organelles in mammalian cells. Biochimica et biophysica acta. 2013;1833:3013–24.

[12] Bharti P, Schliebs W, Schievelbusch T, Neuhaus A, David C, Kock K, et al. PEX14 is required for microtubule-based peroxisome motility in human cells. J Cell Sci. 2011;124:1759–68.

[13] Albertini M, Rehling P, Erdmann R, Girzalsky W, Kiel JAKW, Veenhuis M, et al. Pex14p, a peroxisomal membrane protein binding both receptors of the two PTS-dependent import pathways. Cell. 1997;89:83–92.

[14] Brocard C, Lametschwandtner G, Koudelka R, Hartig A. Pex14p is a member of the protein linkage map of Pex5p. EMBO J. 1997;16:5491–500.

[15] Meinecke M, Cizmowski C, Schliebs W, Kruger V, Beck S, Wagner R, et al. The peroxisomal importomer constitutes a large and highly dynamic pore. Nat Cell Biol. 2010;12:273–7.

[16] Honsho M, Yamashita S, Fujiki Y. Peroxisome homeostasis: Mechanisms of division and selective degradation of peroxisomes in mammals. Biochimica et biophysica acta. 2016;1863:984–91.

[17] Manjithaya R, Nazarko TY, Farre JC, Subramani S. Molecular mechanism and physiological role of pexophagy. FEBS Lett. 2010;584:1367–73.

[18] Nordgren M, Wang B, Apanasets O, Fransen M. Peroxisome degradation in mammals: mechanisms of action, recent advances, and perspectives. Front Physiol. 2013;4:145.

[19] Hara-Kuge S, Fujiki Y. The peroxin Pex14p is involved in LC3-dependent degradation of mammalian peroxisomes. Exp Cell Res. 2008;314:3531–41.

[20] Jiang L, Hara-Kuge S, Yamashita S, Fujiki Y. Peroxin Pex14p is the key component for coordinated autophagic degradation of mammalian peroxisomes by direct binding to LC3-II. Genes Cells. 2015;20:36–49.

[21] Asare A, Levorse J, Fuchs E. Coupling organelle inheritance with mitosis to balance growth and differentiation. Science. 2017;355.

[22] Theiss C, Neuhaus A, Schliebs W, Erdmann R. TubStain: a universal peptide-tool to label microtubules. Histochem Cell Biol. 2012;138:531–40.

[23] Neufeld C, Filipp FV, Simon B, Neuhaus A, Schuller N, David C, et al. Structural basis for competitive interactions of Pex14 with the import receptors Pex5 and Pex19. EMBO J. 2009;28:745–54.

[24] Neuhaus A, Kooshapur H, Wolf J, Meyer NH, Madl T, Saidowsky J, et al. A novel Pex14 protein-interacting site of human Pex5 is critical for matrix protein import into peroxisomes. J Biol Chem. 2014;289:437–48.

[25] Saidowsky J, Dodt G, Kirchberg K, Wegner A, Nastainczyk W, Kunau WH, et al. The di-aromatic pentapeptide repeats of the human peroxisome import receptor PEX5 are separate high affinity binding sites for the peroxisomal membrane protein PEX14. J Biol Chem. 2001;276:34524–9.

[26] Itoh R, Fujiki Y. Functional domains and dynamic assembly of the peroxin Pex14p, the entry site of matrix proteins. J Biol Chem. 2006;281:10196–205.

[27] Aylett CH, Lowe J, Amos LA. New insights into the mechanisms of cytomotive actin and tubulin filaments. Int Rev Cell Mol Biol. 2011;292:1–71.

[28] Mizuno N, Toba S, Edamatsu M, Watai-Nishii J, Hirokawa N, Toyoshima YY, et al. Dynein and kinesin share an overlapping microtubule-binding site. EMBO J. 2004;23:2459–67.

[29] Tomishige M, Vale RD. Controlling kinesin by reversible disulfide cross-linking. Identifying the motility-producing conformational change. J Cell Biol. 2000;151:1081–92.

[30] Larcher JC, Boucher D, Lazereg S, Gros F, Denoulet P. Interaction of kinesin motor domains with alpha- and beta-tubulin subunits at a tau-independent binding site. Regulation by polyglutamylation. J Biol Chem. 1996;271:22117–24.

[31] Woehlke G, Ruby AK, Hart CL, Ly B, Hom-Booher N, Vale RD. Microtubule interaction site of the kinesin motor. Cell. 1997;90:207–16.

[32] Gigant B, Wang W, Dreier B, Jiang Q, Pecqueur L, Pluckthun A, et al. Structure of a kinesin-tubulin complex and implications for kinesin motility. Nature structural & molecular biology. 2013;20:1001–7.

[33] Gouveia AM, Guimaraes CP, Oliveira ME, Sa-Miranda C, Azevedo JE. Insertion of Pex5p into the peroxisomal membrane is cargo protein-dependent. J Biol Chem. 2003;278:4389–92.

[34] Francisco T, Rodrigues TA, Freitas MO, Grou CP, Carvalho AF, Sa-Miranda C, et al. A cargo-centered perspective on the PEX5 receptor-mediated peroxisomal protein import pathway. J Biol Chem. 2013;288:29151–9.

[35] Otera H, Setoguchi K, Hamasaki M, Kumashiro T, Shimizu N, Fujiki Y. Peroxisomal targeting signal receptor Pex5p interacts with cargoes and import machinery components in a spatiotemporally differentiated manner: Conserved Pex5p WXXXF/Y motifs are critical for matrix protein import. Mol Cell Biol. 2002;22:1639–55.

[36] Nguyen T, Bjorkman J, Paton BC, Crane DI. Failure of microtubule-mediated peroxisome division and trafficking in disorders with reduced peroxisome abundance. J Cell Sci. 2006;119:636–45.

[37] Dodt G, Gould SJ. Multiple *PEX* genes are required for proper subcellular distribution and stability of Pex5p, the PTS1 receptor: Evidence that PTS1 protein import is mediated by a cycling receptor. The Journal of cell biology. 1996;135:1763–74.

[38] Skiniotis G, Cochran JC, Muller J, Mandelkow E, Gilbert SP, Hoenger A. Modulation of kinesin binding by the C-termini of tubulin. EMBO J. 2004;23:989–99.

[39] Uchimura S, Oguchi Y, Katsuki M, Usui T, Osada H, Nikawa J, et al. Identification of a strong binding site for kinesin on the microtubule using mutant analysis of tubulin. EMBO J. 2006;25:5932–41.

[40] Barlan K, Gelfand VI. Microtubule-Based Transport and the Distribution, Tethering, and Organization of Organelles. Cold Spring Harbor perspectives in biology. 2017;9.

[41] Halbach A, Landgraf C, Lorenzen S, Rosenkranz K, Volkmer-Engert R, Erdmann R, et al. Targeting of the tail-anchored peroxisomal membrane proteins PEX26 and PEX15 occurs through C-terminal PEX19-binding sites. J Cell Sci. 2006;119:2508–17.

[42] Schliebs W, Saidowsky J, Agianian B, Dodt G, Herberg FW, Kunau WH. Recombinant human peroxisomal targeting signal receptor PEX5. Structural basis for interaction of PEX5 with PEX14. J Biol Chem. 1999;274:5666–73.

[43] Waterman-Storer CM. Microtubule/organelle motility assays. Curr Protoc Cell Biol. 2001;Chapter 13:Unit 13 1.

[44] Merrifield RB. Solid phase peptide synthesis. I. The synthesis of a tetrapeptide. J Am Chem Soc 1963;85:2149–58.

[45] Will GK, Soukupova M, Hong X, Erdmann KS, Kiel JA, Dodt G, et al. Identification and characterization of the human orthologue of yeast Pex14p. Mol Cell Biol. 1999;19:2265–77.

[46] Stanley WA, Filipp FV, Kursula P, Schuller N, Erdmann R, Schliebs W, et al. Recognition of a functional peroxisome type 1 target by the dynamic import receptor Pex5p. Mol Cell. 2006;24:653–63.

[47] Schliebs W, Saidowsky J, Agianian B, Dodt G, Herberg FW, Kunau WH. Recombinant human peroxisomal targeting signal receptor PEX5. Structural basis for interaction of PEX5 with PEX14. J Biol Chem. 1999;274:5666–73.

[48] Theiss C, Napirei M, Meller K. Impairment of anterograde and retrograde neurofilament transport after anti-kinesin and anti-dynein antibody microinjection in chicken dorsal root ganglia. European journal of cell biology. 2005;84:29–43.

[49] Keller S, Vargas C, Zhao H, Piszczek G, Brautigam CA, Schuck P. High-precision isothermal titration calorimetry with automated peak-shape analysis. Anal Chem. 2012;84:5066–73.

[50] Houtman JC, Brown PH, Bowden B, Yamaguchi H, Appella E, Samelson LE, et al. Studying multisite binary and ternary protein interactions by global analysis of isothermal titration calorimetry data in SEDPHAT: application to adaptor protein complexes in cell signaling. Protein Sci. 2007;16:30–42.

[51] Delaglio F, Grzesiek S, Vuister GW, Zhu G, Pfeifer J, Bax A. NMRPipe: a multidimensional spectral processing system based on UNIX pipes. J Biomol NMR. 1995;6:277–93.

